# Gel Electrophoresis/Electroelution Sorting Fractionator combined with Filter Aided Sample preparation (FASP) for deep proteomic analysis

**DOI:** 10.1101/2021.04.16.440150

**Authors:** Yassel Ramos, Alexis Almeida, Jenis Carpio, Arielis Rodríguez-Ulloa, Yasser Perera, Luis J. González, Jacek R. Wiśniewski, Vladimir Besada

## Abstract

Sample preparation and protein fractionation are important issues in proteomic studies in spite of the technological achievements on protein mass spectrometry. Protein extraction procedures strongly affect the performance of fractionation methods by provoking protein dispersion in several fractions. The most notable exception is SDS-PAGE-based protein fractionation due to its extraordinary resolution and the effectiveness of SDS as a solubilizing agent. Its main limitation lies in the poor recovery of the gel-trapped proteins, where protein electro-elution is the most successful approach to overcome this drawback. We created a device to separate complex mixture of proteins and peptides (named “GEES fractionator”) that is based on the continuous *G*el *E*lectrophoresis/*E*lectro-elution *S*orting of these molecules. In an unsupervised process, complex mixtures of proteins or peptides are fractionated into the gel while separated fractions are simultaneously and sequentially electro-eluted to the solution containing wells. The performance of the device was studied for SDS-PAGE-based protein fractionation in terms of reproducibility, protein recovery and loading capacity. In the SDS-free PAGE setup, complex peptide mixtures can also be fractionated. More than 11 700 proteins were identified in the whole-cell lysate of the CaSki cell line by using the GEES fractionator combined with the Filter Aided Sample Preparation (FASP) method and mass spectrometry analysis. GEES-based proteome characterization shows a 1.7 fold increase in the number of identified proteins compared to the unfractionated sample analysis. Proteins involved in the co-regulated transcription activity, as well as cancer related pathways such as apoptosis signaling, P53 and RAS pathways are more represented in the protein identification output of GEES-based fractionation approaches.

## Introduction

Biomolecule separation is mandatory in proteomics. Both, proteins and non-proteinaceous materials are matters of concern during sample preparation. Proteins must be isolated, completely extracted and deeply scrutinized to reach a comprehensive description of the composition of the protein sample, while non-proteinaceous material interfere in most of the procedures and measurements used to address this goal [1].

Sample preparation for two dimensional gel electrophoresis (2DE) was a landmark at the beginning of proteomics by setting procedures that should ensure both, efficient protein extraction and the removal of DNA, lipids, polysaccharides, as well as metabolites [2]. Conventional methods of protein precipitation, delipidation and DNA depletion were re-evaluated and inserted within the new context of sample preparation for 2DE [3, 4]. Although impurities may remain at a low content, an in-gel isolated protein can be considered free of large-size impurities (those not entering or remaining in the gel) or small-size impurities (those readily washed out of the gel) [5]. Therefore, the combination of in-gel protein digestion with mass spectrometry should yield reasonably noiseless spectra. However, this predictable output is frequently limited by the poor peptide recovery of this commonly used method [6]. A similar procedure was later adopted for the direct analysis of SDS-PAGE gel bands, instead of 2DE spots.

Sample preparation in the presence of highly concentrated chaotropic agents or detergent solutions is a common procedure for protein extraction. Typically, detergents must be completely removed, since most of them interfere in downstream steps such as enzymatic hydrolysis, liquid chromatography and mass spectrometry analyses. Often, samples obtained by these methods contain non-proteinaceous molecules that have deleterious effects on chromatography and mass spectra like LC peak widening, ineffective separation, signal suppression effects and a decreased signal to noise ratio. These phenomena greatly contribute to the high ratio of non-assigned MS/MS spectra commonly obtained in LC-MS/MS measurements [7]. To overcome this drawback, sample preparation methods have recently evolved to include the concept of a “proteomic reactor”, which maximize protein and peptide recovery while solubilizing reagents as well as most of the non-proteinaceous impurities are removed in a “one-pot” principle [8]. Notable examples are the Filter Aided Sample Preparation (FASP) [9] and their modified variants such as GOFASP [10] and MED-FASP [11], the single-pot solid-phase-enhanced sample preparation (SP3) [12], in-StageTip (iST) [13], the suspension trapping methods (STRAP) [14] and Protein Aggregation Capture (PAC) [15]. These methods have also been successfully combined with peptide fractionation techniques resulting in a significant increase in the number of identified proteins [16,12,17]. Recently, an average of 11 300 protein groups in 12 cell lines were identified combining the iST method with high pH reversed phase peptide fractionation using the “spider fractionator” device, an automatic rotor-valve-based fraction collector coupled online to the nanoflow HPLC [17].

Sodium dodecyl sulfate (SDS), an anionic detergent, is frequently used as the solubilizing agent. When SDS concentration is higher than its critical micellar concentration, SDS binds to proteins at a w/w ratio of 1.4 g of SDS per gram of protein. Virtually all proteins in a biological sample are considered to be quantitatively extracted at an SDS to protein ratio of 2:1 (w/w). On heating the sample at 95°C in the presence of SDS and reducing agents, there is a denaturing effect that ensures the inactivation of almost all of the enzymatic activity, including the undesirable endogenous protease activity.

The use of SDS in the protein extraction solution was conveniently combined with SDS-PAGE protein fractionation in the GeLC method [18,19]. After protein fractionation by SDS-PAGE, the SDS is removed, the proteins are in-gel digested, and the generated peptide mixture is passively eluted from the gel for further LC-MS/MS analysis. In the GeLC extended variant DF-PAGE method, after the in-gel digestion, the peptide mixture is electro-transferred to a second gel and fractionated by SDS-free PAGE resulting in the increase of the number of identified proteins [20,21]. In this paper, the authors demonstrate the benefits of transferring peptides to a second dimension by using an electric field instead of passive peptide diffusion and loading them into a second gel [20].

The poor recovery of the extracted peptides or proteins from the gel through passive diffusion has limited the use of the gel-based electrophoresis techniques in proteomics in spite of their well-known advantages in terms of resolution and the simplicity of the removal of interfering substances. This led to the successful insertion, in proteomics, of the gel-free electrophoresis techniques, i.e. Free Flow Electrophoresis (FFE) [22], Multi-Compartment Electrolyzer (MCE) [23], as well as the gel-based Off Gel Electrophoresis (OGE) [24] that separates proteins or peptides “in-solution” (FFE and MCE) or in a very low crosslinked gel (OGE). Capillary Electrophoresis (CE) has also received the benefits of improved instrumentation, in both gel-based and gel-free variants, and in particular, the possibility of coupling it directly to the mass spectrometer [25].

Alternatively, electro-elution procedures have been used to recover gel-electrophoresed proteins [26,27,28,29]. In the proteomics context, the GELFREE system is an SDS-PAGE-based protein fractionation device that is frequently used for Top-Down experiments [30,31]. In this horizontally designed device, the gel is in contact with the collection chamber solution that is end-capped with a negatively charged 3.5 kDa molecular weight cut-off membrane. Eluted proteins cannot pass through the membrane and are trapped in the collection chamber. Once proteins are electro-eluted toward the collection chamber, electrophoresis is halted and the fraction is collected. Before restarting the electrophoresis, the collection chamber should be washed and refilled for the electro-elution of the next fraction in the same chamber [27].

More recently, the SDS-free PAGE technique was adapted to an on-line electro-elution system named Gel Electrophoresis/Electroelution Sorting (GEES) [32]. The system is based on the original “Disc Electrophoresis” carried out in tubes. The bottom of the gel tube is in contact with the collection unit solution and the eluted peptides are retained by a 0.5 kDa molecular weight cut-off membrane. When the peptide fraction is collected, the gel tube and cathode chamber are manually positioned at a new collection unit to electro-elute the next fraction [32]. Here, we present a new GEES-based device for both, protein and peptide fractionation (GEES fractionator). The equipment permits, in an unsupervised process, the simultaneous in-gel protein or peptide separation and the in-solution collection of the fractionated biomolecules. The performance of the device is discussed in terms of reproducibility, recovery and loading capacity. The GEES fractionator for protein SDS-PAGE or peptide SDS-free-PAGE fractionation combined with the MED-FASP method enabled the identification of more than 11 700 proteins in the CaSki cell line.

## Materials and Methods

### Materials and Reagents

All reagents for gel electrophoresis were obtained from Bio-Rad (USA). The Low Molecular Weight Markers were also provided by Bio-Rad. The 0.5 kDa and 3.5 kDa MWCO dialysis membranes were purchased from Spectrum Laboratories (USA). Tris soluble proteins from the *Escherichia coli* strain W3110 cells were obtained by sequential solubilization as described by Molloy *et al* [33]. The CaSki human cell line was originally acquired from American Type Culture Collection (Rockville, MD, USA).

### Sample Preparation

Standard protein solutions, *E. coli* proteins as well as CaSki cells were incubated for 5 min at 95°C in the electrophoresis sample buffer (1% SDS, 2.5% β-mercaptoetanol, 12.5% glycerol, 62.5 mM Tris/HCl, pH 6.8). CaSki cell extraction was performed by sonication. For cysteine modification, acrylamide was added to 6.25% (w/v) and the samples were incubated for 1h at 25°C. Insoluble material was removed by centrifugation at 16,000 x g, 20°C for 20 min. Protein and peptide concentration was estimated by a 96-Well-Plate-Based Tryptophan Fluorescent Assay as described previously [34].

### GEES equipment

For a better description of this equipment, four parts or systems will be considered below: electrophoresis system, mechanical system, electronic system and software.

*The electrophoresis system* has five components: a cathode chamber, tubes with the polyacrylamide gel column, the collection device, the anode chamber and the power supply (Figure 1, Supplementary Information I). The cathode chamber is attached to a mechanical arm and contains the cathode electrode. There are 4-8 holes in its underside where the tubes containing the gel (od 5 mm, id 3 mm, length 125 mm) are inserted. The upper ends of the tubes are open for the contact of gel with the cathode buffer solution and for sample loading. The bottom ends of the tubes are open allowing the gel contact with the collection wells solution while electrophoresis is taking place. The collection device is a 96-well format plate. The bottom of the wells consists of a replaceable membrane (0.5 kDa or 3.5 kDa) that holds the collection buffer solution (on the upper side) and makes contact with the anode buffer solution (on the bottom side). The anode chamber contains the anode electrode. The power supply controls the current for electrophoresis.

**Figure 1.**
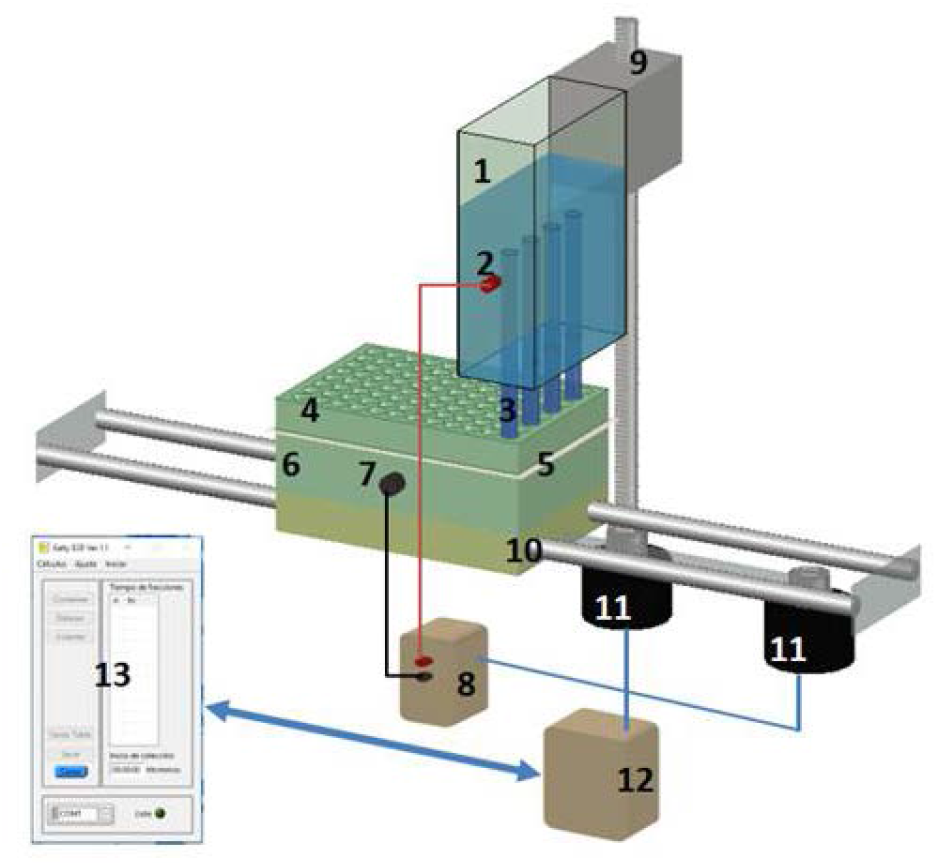
Schematic representation of the GEES Fractionator. Cathode chamber (1), access to the cathode electrode (2), tubes containing gel (3), 96-well format collection tray (4), membrane for peptide or protein trapping (5), anode chamber (6), access to the anode electrode (7), power supply (8) bracket with endless screw for vertical movement (9), holder with two rail for horizontal movement (10) Stepper motors (11), electronic circuits unit (12), GEES software (13).

*The mechanical system* consists of: a bracket with an endless screw that enables the vertical movement of the cathode chamber and the gel containing tubes; a holder with two rails that enables the horizontal movement of the collection unit and two stepper motors in charge of the vertical and horizontal movements of the cathode and anode chambers, respectively (Figure 1, Supplementary Information I).

*The electronic system* is controlled in a synchronized way, by the GEES software. It is made up of a set of sensors, electronic circuits and two stepper motors. Through them, the horizontal and vertical movements of the electrophoresis chambers are achieved and, in addition, the power supply is enabled and disabled each time the electrophoresis is interrupted for the samples collection.

*The GEES software* uses the Windows (XP and more recent versions) operating system. It was developed with the Laboratory Virtual Instrument Engineering Workbench (Labview 2009) platform (National Instrument, www.ni.com). The software controls timing and positioning (Supplementary Information I). There are four collection methods:

1. Automatic – using a semi-empirical formula [32], the software calculates the collection time for each fraction having the following input data: 1) the time taken for the front of the dye to reach the bottom end of the tube and 2) the number of fractions that will be collected. The software timer automatically turns on when the electrophoresis starts, and it gives the time value.
2. Semiautomatic – using a semi-empirical formula [32], the software calculates the collection time for each fraction having the following input data: 1) the time introduced by the user and 2) the number of fractions that will be collected.
3. Custom time/fraction – the user introduces the collection time for each fraction and the number of fractions that will be collected.
4. Common time/fraction – the user introduces a single value for collection time that is equal for all fractions, as well as the number of fractions that will be collected.

### GEES operating conditions

The length of both, the stacking and resolving gels depend on the user’s choices. Unless otherwise noted, the gel column contains a 5.6 cm long resolving gel and a 2.8 cm long stacking gel. Gels, sample buffer and running buffer were prepared for SDS-PAGE protein fractionation as described by Laemmli [35]. The resolving and staking gels were polymerized at 15% T, 3% C and 6%T, 3%C respectively. For SDS-free PAGE peptide fractionation, all buffers and gels composition were identical but excluding SDS. The electrophoresis was performed at 2 mA per gel. The collection wells were bottom-capped with a 0.5 kDa (for peptides) or a 3.5 kDa (for proteins) molecular weight cut-off membrane and provided with 200 µL of running buffer. Since protein samples were prepared in the electrophoresis sample buffer, only bromophenol blue traces were added before loading. For peptide fractionation, the peptide mixture was diluted twice in glycerol 25%, Tris/HCl 125 mM, pH 6.8.

### Analytical SDS-PAGE

Unfractionated samples and GEES protein fractions were analyzed by discontinuous SDS-PAGE using 15% T resolving gels as described by Laemmli [35]. Equivalent volumes of the fractions were diluted with equal volumes of the sample buffer 2X (SDS 2%, β-mercaptoethanol 5%, glycerol 25%, Tris/HCl 125 mM, pH 6.8 supplied with traces of bromophenol blue) and loaded into the individual slab gel lanes. Proteins were silver stained in-gel according to Heukeshoven and Dernick [36].

### Proteomic sample preparation

Unfractionated samples (80 µg) and GEES protein fractions (2-16 µg) were processed by the MED-FASP protocol using Microcon 30k centrifugal ultrafiltration units (Merck, Darmstadt) [11]. Samples were mixed with 250 µL of the UA buffer (8 M urea, 100 mM Tris/HCl pH 8.5), loaded into the filter unit and centrifuged at 10,000 x g at 20°C for 20 min. The eluates were discarded, 250 µL of UA was again pipetted into the filter unit, the units were centrifuged and the eluates discarded. This step was repeated 5 times followed by 3 washings with the DB buffer (50 mM Tris/HCl pH 8.5). Proteins were digested in 80 µL of the DB buffer at 37°C for 18 h, using endoproteinase LysC at an enzyme to protein ratio of 1:40 (w/w). The peptides released were collected by centrifugation at 10,000 × g for 10 min followed by two washings with 125 µL and 100 µL of DB respectively. The material remaining on the filter was digested with 1.5 µg of trypsin in the DB buffer at 37°C for 6 h. The peptides released were collected as described for LysC digestions. A volume of 8 µg of total peptide of unfractionated samples or 2-5 µg of GEES protein fractions were desalted and stored at 4°C.

### LC-MS/MS and data analysis

Liquid chromatography was performed on a Proxeon Easy-nLC System (Thermo Fisher Scientific). Peptide separation was carried out at 300 nL/min for 120 min on a 50 cm C18-column using an acetonitrile gradient of 5-30% (v/v) in 0.1% (v/v) formic acid. The columns were thermostated at 60°C. Analysis of the on-line eluted peptides was performed using a QExactive HF-X mass spectrometer (Thermo-Fisher Scientific). The mass spectrometer operated in a data dependent mode with survey scans acquired at a resolution of 60,000 at m/z 400. Selection for fragmentation consisted of the 15 most abundant precursor ions from the survey scan with a charge ≥ +2 within the 300-1 650 m/z range. All MS/MS spectra were searched using the MaxQuant software (v 1.6.2.10) [37]. Propionamide-cysteine was set as a fixed modification, whereas oxidized methionine (sulfoxide), acetylated N-terminal protein and the deamidation of asparagine and glutamine were set as variable modifications. Minimum peptide length was specified as 7 amino acids. The first peptide tolerance search was set at 20 ppm, while the main peptide tolerance search was at 4.5 ppm. The maximum false discovery rate for peptides and proteins was specified as 0.01.

### Bioinformatic tools

The functional classification of proteins was based on the information annotated in the Gene Ontology (GO) database (http://www.geneontology.org). The analysis was performed using the Panther Classification System (http://pantherdb.org) [38]. Proteins were also classified according to biological pathways reported in the Panther database [39]. The human genome was the reference background for the analysis.

## Results and Discussion

### GEES electrophoresis unit design

The GEES fractionator uses a vertical tube-based electrophoresis (figure 1, Supplementary Information I). The user can select the resolving gel length, depending on the resolution and the mass range separation required. Shorter and lower concentration crosslinked resolving gels makes it possible for high MW protein to elute in a reasonable experimental time. Longer gel column generally increase the resolution of low to medium MW proteins (10-60 kDa), but they require a proportionally longer separation time. This can affect the resolution by widening the band through diffusion during their migration along the resolving gel in the long electrophoresis period, mainly for high MW proteins. A satisfactory balance should be tested for specific aims. For standard proteomic studies (protein fractionation of a proteome), we set our experimental conditions for the best fractionation performance for 3-150 kDa MW proteins by using a 15%T, 5.6 cm resolving gel length. Since with this equipment sample collection does not need to be supervised, long electrophoretic runs can be easily carried out. Consequently the relatively simple high MW subproteome (ex: > 150 kDa) can be collected in few fractions, or in only one fraction, during a long collection period.

Together with the vertical and horizontal movement of the cathode and anode respectively, the vertical tube contacts the solution of independent collection wells (figure 1, Supplementary Information I) along the electrophoretic run. Thus, neither the electrophoresis halting/running steps nor the washing and refilling steps of the collection chambers, are required. The collection of protein or peptide fractions in individual wells also helps to diminish cross contamination among consecutive fractions.

The collection unit is a 96-well format plate making it compatible with most of the instrumental formats, including the available nanoHPLC autosampler tray (figure 1, Supplementary Information I). The bottom of the wells is capped with a membrane for protein (3.5 kDa cut-off) or peptide (0.5 kDa cut-off) trapping. Usually, 200-300 µL of the collection solution is used in all experiments, and no dilution is therefore obtained, since several hundred microliters of the unfractionated sample can be loaded into the system.

The four modes used to schedule collection time, offer a range of possibilities to the user for different applications. For proteome fractionation, the automatic collection mode is usually used, since it offers similar partitioned fractions in terms of the amount of protein, which is frequently desired for increasing the number of identified proteins in bottom-up proteomics. For a particular protein enrichment, the *Custom time/fraction* collection is the mode of choice. Equal collection time for all fractions can also be set using the *Common time/fraction* collection mode.

### Reproducibility

Two samples were used for the reproducibility evaluation: the Bio-Rad Low Molecular Weight Markers (MWM) and the *E. coli* proteome. For the MWM, a preliminary experiment using the automatic collection mode of the GEES fractionator allows us to predict the collection time of the five lowest molecular weight proteins [lysozyme (14 kDa), soybean trypsin inhibitor (21 kDa), carbonic anhydrase (31 kDa), ovalbumin (45 kDa) and bovine serum albumin (66 kDa)]. We can therefore collect them in subsequent short-time experiments using the *Custom time/fraction* collection mode (the collection time per well is pre-defined by the user). Using the new parameters, three MWM protein fractionation experiments were performed in an unsupervised manner and on different days (using only one channel and the same stock solutions). The first 8 fractions were analyzed by SDS-PAGE and the reproducibility criterion was to detect the protein markers in the same and single fraction. Figure 2A shows that the MWM bands were always detected in the same fraction.

**Figure 2.**
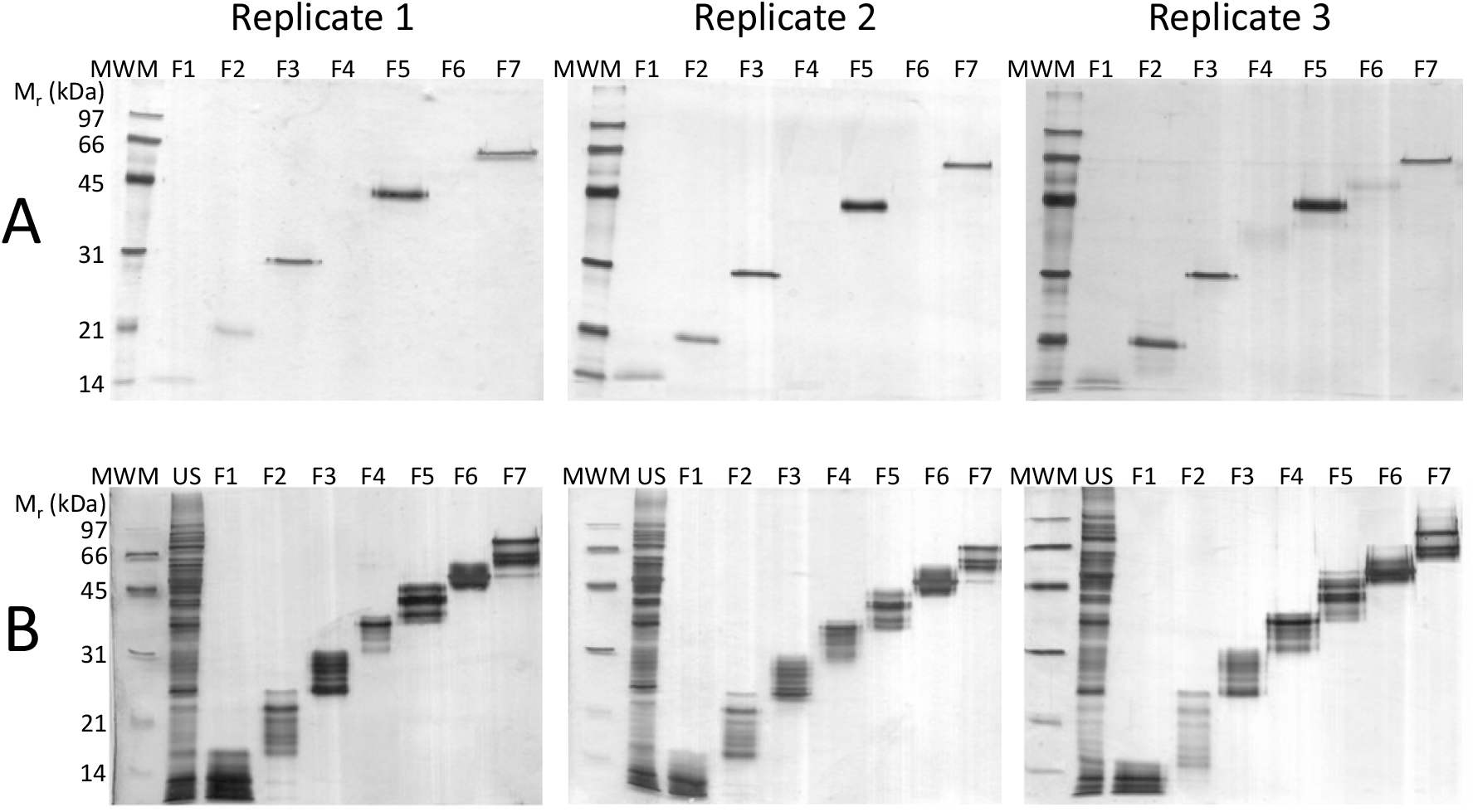
Reproducibility achieved with the SDS-PAGE-based GEES fractionator with automatic or programmed collection. A and B are analytical SDS-PAGE of the collected fractions (A): Three replicates of the Bio-Rad Low Molecular Weight Marker fractionation carried out on different days. (B): Three replicates of the *E. coli* Tris soluble protein fractionation carried out on different days. MWM: Molecular Weight Marker. US: Unfractionated sample.

For *E. coli* proteome fractionation, the GEES fractionator was used in the automatic collection mode. The band profiles in all fractions were similar in all experiments carried out on different days. Very few bands migrating at the boundaries of the band profiles were detected in consecutive fractions, or in only one of the consecutive fractions (figure 2B). In our hands, reproducibility depends highly on the consistency of the solutions and gel casting in terms of composition, ionic strength and pH of the solutions, as well as gel column length. The collection process starting time is also a crucial variable. It is defined by the arrival of the dye front to the bottom end of the tube. The more efficient the stacking process is, the narrowest the front of the dye; thereby resulting in a higher precision of the collection starting time. Controlling these operation variables, the GEES fractionator provides highly reproducible protein fractionation as shown in figure 2B.

### Protein recovery

Protein recovery is affected by several factors including physicochemical protein properties, residence time and temperature in the collection well, the amount of fractionated proteins, absorption properties of the container polymers and membrane, among others. All these factors are protein dependent, including those related to the container properties, therefore, values of protein recovery should be considered with caution. Residence time in the collection well, for example, is protein size-dependent. Collection time for small proteins should be shorter than for larger proteins. Even for an individual well, residence time of the first eluted proteins is similar to the collection time of the entire fraction, whereas it varies to a few seconds for the last proteins eluted in the same well. Higher protein loses are usually obtained for highly hydrophobic and larger proteins than for hydrophilic and small proteins, although the presence of SDS tends to reduce this drawback. Accordingly, we estimated protein recovery of mixtures instead of individual proteins in order to average all these factors.

For the six MWM proteins with a molecular mass ranging from 14 to 97 kDa and 500 ng each, protein recovery was estimated by 280 nm absorbance of the combined fractions compared to the unfractionated sample. An average protein recovery of 83% was calculated from three different GEES protein fractionations carried out on different days (Supplementary Information IIA). For a cell line with a complex protein extract containing a wide range of protein abundance and molecular weight proteins, the typical protein recovery achieved is about 75% (Supplementary Information IIB).

### Loading capacity

The GEES fractionator prototype uses a 3 mm id tube. Loading capacity was studied for this size tube using the automatic collection mode with an increasing protein amount of *E. coli* proteome ranging from 0.25 to 1.5 mg. The corresponding volume ranged from 50 to 200 µL (for 1.5 mg of the loaded sample (300 µL), the sample was concentrated to 100 µL by vacuum). Figure 3A-C shows the SDS-PAGE analysis of the 8 fractions collected automatically during the first 5 h. High quality fractionation was obtained among all the range of protein loading studied (figure 3A-C).

**Figure 3.**
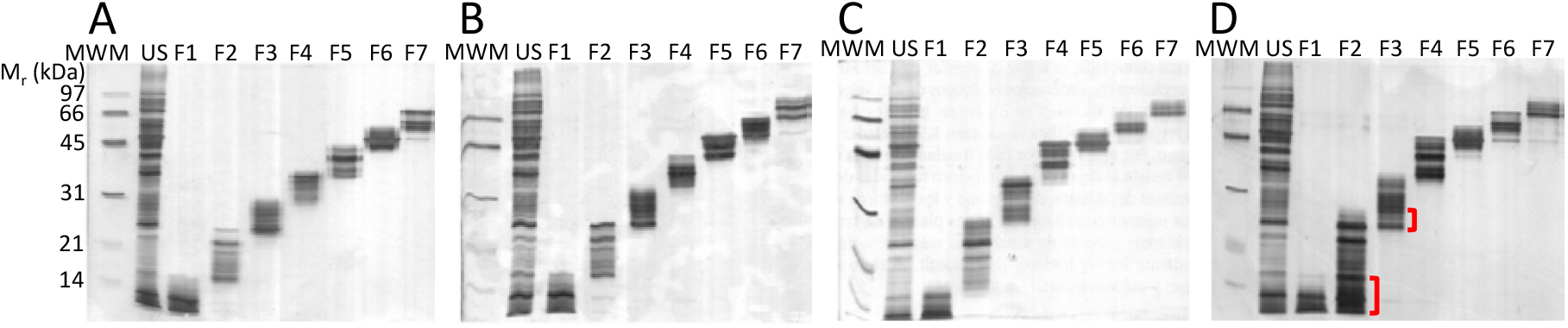
Loading capacity of the SDS-PAGE-based GEES fractionator. Fractions F1-F7 of the *E. coli* Tris soluble protein GEES fractionation, 250 µg in 50 µL (A), 1 mg in 200 µL (B), 1.5 mg in 100 µL (C), 1.5 mg in 300 µL (D). Brackets indicate protein dispersion among fractions F1 and F2. MWM: Molecular Weight Marker. US: Unfractionated sample.

Particular precaution should be taken with loading volume. Since loading volume is not defined by the device, but by the user, the loading-to-stacking gel volume ratio should be established for an efficient sample stacking before protein separation. Figure 3D shows the results obtained for 1.5 mg of sample loading (300 µL of the total loading volume) without prior vacuum concentration. A strong overlapping between fraction 1 and 2 was obtained (brackets in figure 3D). The poor resolution observed, mainly for the low molecular weight fractions, suggests that there is incomplete sample stacking due to the high sample volume loaded. Although the main factor for sample stacking is the discontinuous buffer system working under the conditions of the Kohlrausch “regulating function” [40], the abrupt change in porosity from the stacking to the resolving gel (6%T to 15%T) also affects protein band stacking. This effect is more pronounced for medium and high molecular weight proteins and it is less significant or even negligible for very low molecular weight proteins (≤ 10 kDa), consequently the deteriorated resolution is not observed for fractions 3-8.

Ever since the stacking gel was introduced by Ornstein [41], the PAGE techniques resolution has improved considerably. Proteins can be concentrated more than 10 000 times during the stacking process, consequently migrating bands of few millimeters wide (sometimes more than 1 cm) are reduced to hundreds of microns just before arriving at the separating gel. The resolution of SDS-PAGE depends largely on the quality of the stacking process. For the GELFREE system, for example, a long stacking gel was used for an efficient protein stacking and a relatively short resolving gel for the electro-elution of high molecular weight proteins in a reasonable experimental time. For the GEES fractionator prototype we used a 2.8 cm length stacking gel, therefore a loading volume of up to 200 µL (2.8 cm length for a 3 mm id tube) can be loaded without affecting resolution. Using these settings, high quality protein fractionation can be obtained for at least up to 1.5 mg of protein loading.

### GEES Protein fractionation for Bottom-Up proteomics

Protein fractions obtained after the SDS-PAGE-based GEES fractionation contain SDS, glycine and the Tris/HCl buffer. The FASP sample preparation method has demonstrated that it can efficiently remove SDS as well as other impurities from the sample [42,43]. The multienzyme digestion variant (MED)-FASP increases the number of identified proteins and their sequence coverage [11]. Thus, applying the MED-FASP method to the samples previously fractionated by SDS-PAGE, using the GEES fractionator, provides a fully compatible straightforward combination for Bottom-Up proteomic experiments.

An SDS 150 µg protein extract from the CaSki cell line was loaded into the GEES fractionator and twelve fractions were collected by the automatic mode. Both, the unfractionated sample and the GEES protein fractions were processed by the MED-FASP method and analyzed by LC-MS/MS. Raw individual files were submitted to protein identification using the MaxQuant software. Combining the 12 MaxQuant output, a total of 91 390 unique peptides corresponding to 10 212 protein groups were identified. The number of proteins identified per fraction ranged from 2 386 to 3 926 protein groups (Supplementary Information III). Although the analysis of the fractions by SDS-PAGE/silver stain shows excellent resolution (figure 4A), 49% of proteins were detected in more than two fractions, probably due to cellular protein processing or protein degradation, as well as protein dispersion during fractionation. Even so, proteome coverage increased in more than 50% compared to the unfractionated protein analysis (6 789 protein groups, Supplementary Information IV) using the same LC-MS/MS settings.

**Figure 4.**
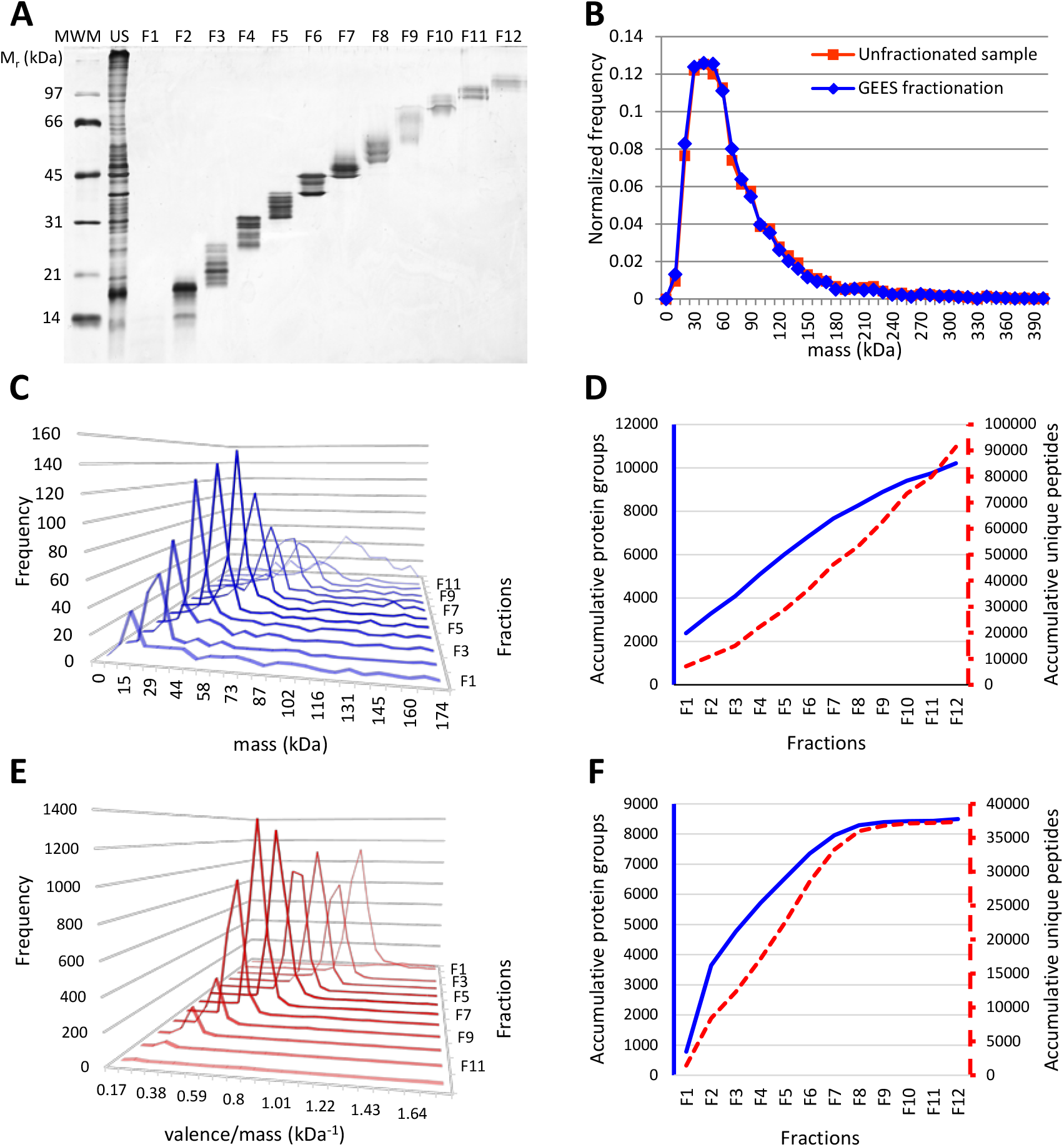
GEES fractionator-based analysis of the CaSki proteome. (A): SDS-PAGE/silver stain analysis of the protein fractions obtained after the SDS-PAGE-based GEES fractionation of 150 µg of the CaSki proteome. F1-12: fractions 1-12 (a volume corresponding to 1/25 of the fraction volume was analyzed), MWM: Molecular Weight Markers. (B): Normalized Molecular Weight distribution of the identified proteins using the GEES fractionator/MED-FASP approach (blue) or the MED-FASP method of the unfractionated sample (red). (C): Frequency per fraction of the protein molecular mass identified in only one fraction (post-translational modifications were not considered for protein mass calculation). (D): Cumulative identified proteins (blue) and peptides (red) counts for the GEES fractionator/MED-FASP approach. (E): Frequency per fraction of the peptide valence to mass ratio, of the unmodified peptides identified in only one fraction. (F): Cumulative identified proteins (blue) and peptides (red) counts for the FASP/GEES fractionator approach.

Three hundred seventy-seven proteins with molecular weights of over 200 kDa were identified (3.7% of all identified proteins). Titin (MW 3 816 kDa) was the identified protein with the highest molecular weight. Processed or partially degradated forms of titin and others very high molecular weight proteins were identified in this experiment (titin was identified in fractions 12-11 and 5-4) since these high molecular weight proteins do not enter the gel in their entire native form.

The identified protein with the lowest molecular weight was the CKLF-like MARVEL transmembrane domain-containing protein 7 (4.5 kDa). In the unfractionated sample, the molecular weight range of the identified proteins was similar; i.e. Titin (MW: 3 816 kDa) was the largest protein and the Translin-associated factor X-interacting protein 1 (4.0 kDa) was the smallest protein. A slightly higher proportion of high molecular weight proteins (molecular weight higher than 200 kDa) was identified in the unfractionated sample (307 proteins corresponding to 4.5% of the total identified proteins). However, the molecular weight mass of the proteins identified with the SDS-PAGE-based GEES fractionator had a similar distribution to that of the unfractionated proteins (figure 4B). Varying the porosity and length of the resolving gel, as well as increasing the collection time, could increase the visibility of the high molecular weight subproteome.

The frequencies of the theoretical molecular weight of the proteins identified in only one fraction (regardless of the post-translational modifications) were plotted against the number of the fraction where they were identified. Figure 4C shows the expected molecular weight dependent fractionation profile for the SDS-PAGE-based methods. Our results confirm the advantage of using fractionation methods at the protein level in proteomics, since any fraction contribute to the total number of identified proteins in a cumulative protein count analysis (figure 4D). The relatively limited use of protein fractionation methods in shotgun proteomics is probably due to the large range of protein solubility, molecular mass, charge and hydrophobicity that the separation techniques have to deal with. Compared to peptide fractionation techniques, the wider range of physicochemical properties of proteins is a challenge for any protein fractionation method as well as making them compatible with later steps. SDS-PAGE is perhaps the only technique that can overcome most of these drawbacks in a unique, simple, and robust way, and with high resolution. Additionally high molecular weight contaminants such as DNA do not enter the stacking and/or the resolving gel, neither do the low molecular weight contaminants such as lipids, glycosides and metabolites, which are removed within the front of the dye. The poor protein recovery of the gel-trapped proteins and the presence of SDS, both considered to be the two main disadvantage of this technique, are solved here by combining the GEES fractionator and the FASP methods.

### GEES peptide fractionation for Bottom-up proteomics

SDS-free PAGE for the fractionation of complex peptide mixtures was introduced several years ago as the second fractionation step of the DF-PAGE method [20]. In this system, only negatively charged peptides at buffer sample pH can migrate through the gel and separate according to their charge and mass. In the discontinuous buffer system Tris/Glycine designed by Ornstein [41], the pH of the sample buffer is 6.8. Accordingly, SDS-free PAGE reduces the complexity of the peptide mixture since only peptides with pI<6.8 are selected for separation and analysis. The human proteome has an average of 16 peptides per protein with pI<6.8 and more than 99.8% of the proteins contain at least one tryptic peptide at this pI range [44].

For peptide fractionation using the SDS-free PAGE-based GEES fractionator, only two settings are different from protein fractionation: 1) the SDS is absent in all the electrophoretic buffers (cathode, sample, stacking gel, resolving gel, collection and anode buffer) and 2) the collection well contains a 0.5 kDa cut-off membrane for peptide retention.

For this approach, CasKi cell proteins were extracted in a SDS containing buffer and they were processed by FASP. The resulting peptide mixture (obtained in the SDS-free Tris/HCl buffer) was diluted in the SDS-free sample buffer and divided into 12 fractions using the automatic collection mode of the GEES fractionator. Peptide mixtures were desalted and analyzed by LC-MS/MS.

In the absence of SDS, peptides migrate according to their charge/size ratio. In practice, the molecular mass of the peptide and the calculated peptide valence are frequently used instead of peptide size and charge, respectively. Considering the unmodified peptides that were identified in only one fraction, the peptide fractionation profile shows a clear dependence of peptide valence/mass ratio (z/m) with electrophoretic migration (figure 4E). Other equations for the dependence of electrophoretic peptide mobility with peptide valence and molecular mass have been described [45]. Experimental data from paper and capillary electrophoresis show that electrophoretic peptide mobility is proportional to z/m^2/3^ (the Offord model [46]) or ln(Z+1)/m^s^, where ***s*** has values between 1/3 and 2/3 depending on peptide properties [45]. The ln(Z+1)/m^2/3^ peptide values calculated from our data against the fraction number show a similar profile to the calculated z/m values shown in figure 4E (data not shown). Similar relationship was previously reported for SDS-free PAGE peptide fractionation by using GEES fractionation with the manual format [32].

Combining the 12 peptide fraction analyses, 37 338 unique peptides corresponding to 8 492 protein groups were identified. The relatively lower peptide to protein ratio (4.4 peptides/protein) reflects the reduced complexity in the peptide mixture as previously described for the SDS-free PAGE technique [44]. An increase of the number of identified protein groups of more than 25% was obtained compared to the unfractionated sample, while 1141 protein groups were identified by this approach, alone. The number of peptides detected in fractions 10-12 decreases significantly, and the number of newly identified proteins from fractions 8 to 12 is low in the cumulative count of identified proteins (figure 4F). However, in two out of the 12 fractions (Fractions 6 and 7), the number of identified proteins is higher than 4200 protein groups (Supplementary Information V). This is higher than the largest number of proteins identified in one fraction of the protein SDS-PAGE-based GEES fractionation strategy (3926 protein groups in fraction 12, Supplementary Information III). These results suggest that the automatic collection mode set for the protein fractionation-based GEES fractionator could be readjusted for peptide fractionation in order to obtain a more uniformly distributed number of peptides per fraction. The user can easily set this parameter, depending on the features and complexity of the peptide mixture.

### CaSki proteome

The CaSki cell line is a human cervical cancer cell line that is widely used in cancer research. Several 2DE-proteomics-based studies searching for biomarkers, as well as studies describing the effect of cancer drugs have used the CaSki cell line [47,48,49]. However, there are few reports on the comprehensive characterization of the proteome of this HPV-16 positive human carcinoma compared to the in-depth proteome characterization of Hela, an HPV-18 positive human cervical carcinoma cell line [11]. Here we report a more in-depth access to the CaSki proteome using the GEES fractionator. On considering the results obtained from both, GEES protein fractionation/MED-FASP and FASP/GEES peptide fractionation approaches, as well as the unfractionated sample analysis, we were able to identify 11 702 protein groups and 127 972 unique peptides (figure 5A). To our knowledge, this is the largest proteome ever described for the CaSki cell line.

**Figure 5.**
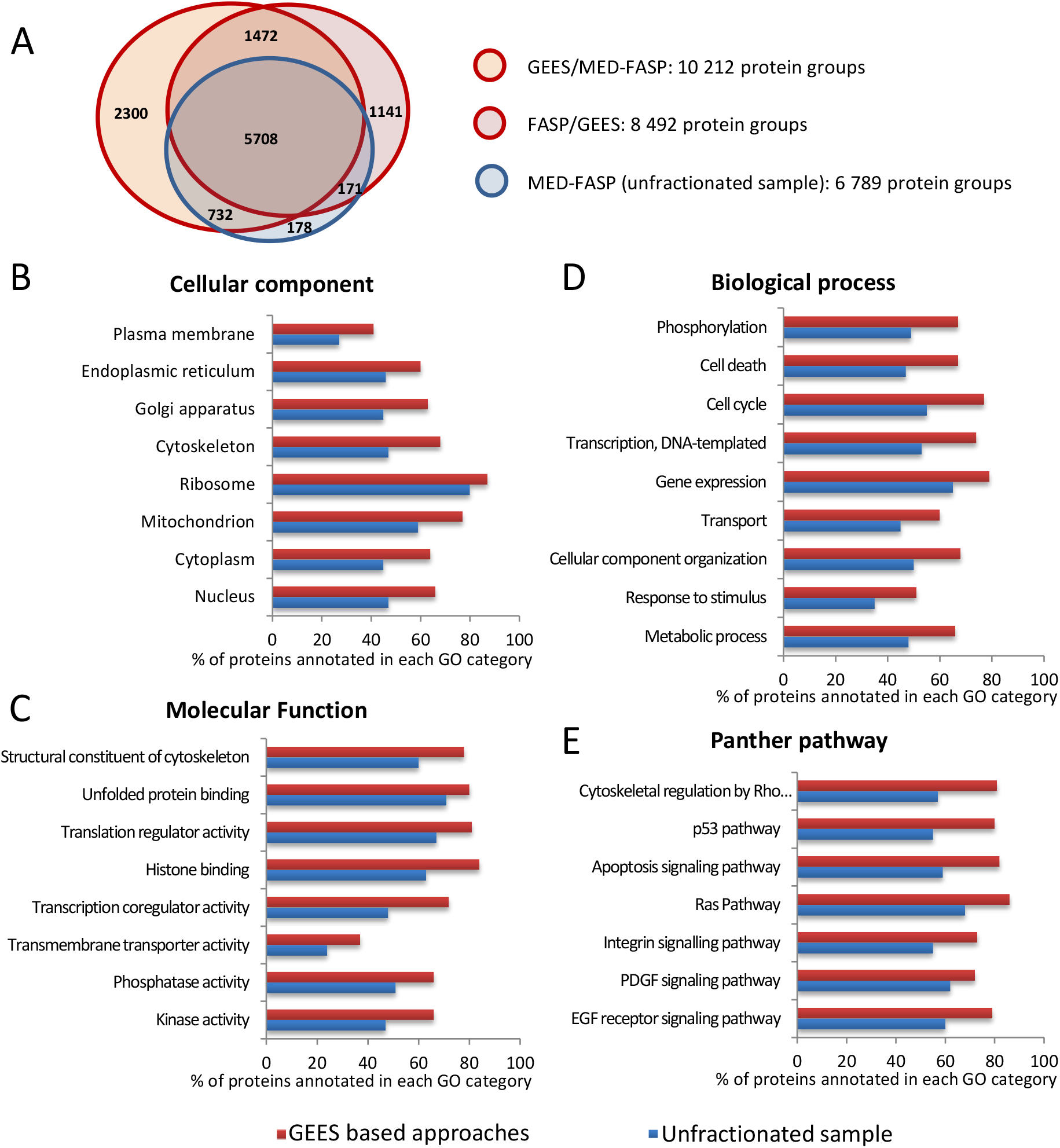
Gene Ontology annotations of the identified proteins: Venn diagram of the number of identified proteins per method (A), cellular component (B), molecular function (C), biological process (D), and Panther pathways (E) of proteins identified without fractionation (blue bars) and after GEES fractionation (red bars) of CaSki total cell lysate. Bars show the percentage of identified proteins with the indicated annotation in relation to the total number of proteins annotated on such biological category.

The combination of the GEES-based protein fractionation method with the MED-FASP method was found to yield the largest number of identified proteins (10 212 protein groups), with 2 300 protein groups only identified by this approach (figure 5A). However, 1141 and 178 protein groups were uniquely identified by the FASP-GEES based peptide fractionation approach and the unfractionated sample (MED-FASP) respectively (figure 5A). Although a more realistic comparison should have a replicate containing experiments with several cell lines or biological tissues, this result confirms the complementary effects usually obtained in the proteomics methodological platform.

Different to other pre-fractionation instruments, the GEES fractionator allows separating both, protein and peptide as independent approaches or combined (i. e. via DF-PAGE method [20]). A balance between the number of generated fractions and the measuring time available for mass spectrometer analysis as well as the starting material amount frequently define the strategy to follow and consequently the proteome depth to reach. For example, by using the “spider” instrument, a peptide fractionator that minimize sample losses, the authors identified 10 769 proteins groups in the 24-fractions instrument format and 60 µg of HeLa protein extract (without using *match between runs* MaxQuant tool) [17]. Scaling down the number of fractions and the starting material to 4 and 20 µg respectively, around 90% of the proteins identified in 24 fractions are still identified, although the number of assigned peptide had a more pronounced decreasing (62%). In our case after fractionating 150 µg of CaSki protein extract at protein and peptide level independently by means of the GEES fractionator, which translate to 24 fractions (12 each one), 11 524 protein groups were identified (11 702 protein groups if the un-fractionated sample analysis and a total of 25 LC-MS/MS measurement are considered). If only GEES-based protein fractionation is considered (meaning 12 fractions), 10 212 protein groups are identified, confirming the benefit of fractionating at protein level.

In addition, we analyzed the effect of the GEES fractionator-based approach on the increase of proteome accessibility in terms of molecular function, biological processes, metabolic pathways and cellular components of the identified proteins. The Gene Ontology terms and the biological pathways represented on both datasets (GEES fractionated and unfractionated samples) were computed using the Panther Classification System. As a reference set, the human genome was used to calculate the percentage of identified proteins in each functional category (figure 5B-E).

While the representativeness of cellular components constituted by relatively highly abundant proteins such as ribosome shows a moderate increase in the GEES-based method, as compared to the unfractionated sample (86.7% vs 80.4% respectively, figure 5B), a considerable increase (with a range of 13-22%) was observed for the rest of the cell components (figure 5B). A similar significant increase (with a range of 10-24%) was obtained for all the molecular functions analyzed. Interestingly, an increase of 23.7% (71.7% vs 48%) was observed in the proteins involved in the co-regulated transcription activity (figure 5C). In relation to the biological process and Panther pathways included in our analysis (figure 5D-E), the GEES fractionator allowed us to access 15-21% and 10-25% more proteins, respectively, compared to the unfractionated sample in all categories. For the proteins related to the p53 pathway, a critical pathway in cancer diseases, we were able to achieve a 25.3 % higher representation in the protein identification output of the GEES fractionated methods.

The MS signal intensity of identified peptides can be used for the quantitative estimation of cellular protein composition and cell size using the Total Protein Approach [50] and the “proteomic ruler” [51]. Our MS data shows that histones intensity represent 2.45% of the total protein intensity of the CaSki protein digestion. This value lies within the range of the histones content (2-4%) reported for different human cell lines (i.e. A549, Hep–G2, PC–3 and U87–MG cell lines) [51]. Using histones as the “proteomic ruler”, the CaSki cellular protein mass was estimated in 266 pg/cell. The 41 and 689 most abundant proteins represent the 25% and the 75% of the total cellular protein mass respectively. These values are similar to those estimated for Hela cells (40 and 600 most abundant proteins respectively), a human cell line that share a common epithelial cervix cancer origin [52].

Typically, the total cellular protein concentration is reported in a range of 20-30% (i.e. 200 to 300 pg/pL) for many cell types [53]. This value can be used to translate the cellular protein mass to cell size. Assuming a spherical cell shape and selecting the medium value 250 pg/pL as the total cellular protein concentration, the estimated diameter of the CaSki cell is 12.7 µm (or a range from 11.9 to 13.6 µm considering the extreme values 200 and 300 pg/pL as the total cellular protein concentration). Although cell size can vary with the cell cycle progression and the culture conditions, these values remarkably match with the reported CaSki cell diameter of 12.3 µm estimated by flow cytometry [54].

The copy number of the identified proteins per cell was also calculated using histones as the “proteomic ruler”. The value of the histone fraction stabilizes from around 12,000 identified peptides or more [51]. This proteome depth is easily attainable with many experimental setups. However, to access the low abundant protein identification and therefore accurately estimate the copy number of the proteins belonging to this sub-proteome, sample pre-fractionation steps are frequently needed. In any scenery, several obligate protein complexes can be used as a “quality control” of the copy number estimations as they are well characterized in terms of their stoichiometric composition. The figure 6A shows our “proteomic ruler” derived copy numbers of the TRiC chaperone complex (T-complex protein 1). Close to the expected 1:1 stoichiometry is obtained among all subunits in the analysis of both, the un-fractionated sample and the GEES-based fractionated sample data (Figure 6A). Additionally, the calculated copy number of the TRiC members for CaSki cells (3-9 x 10^6^ copy number) is within the range of those obtained for other human cell line reported previously [51].

**Figure 6.**
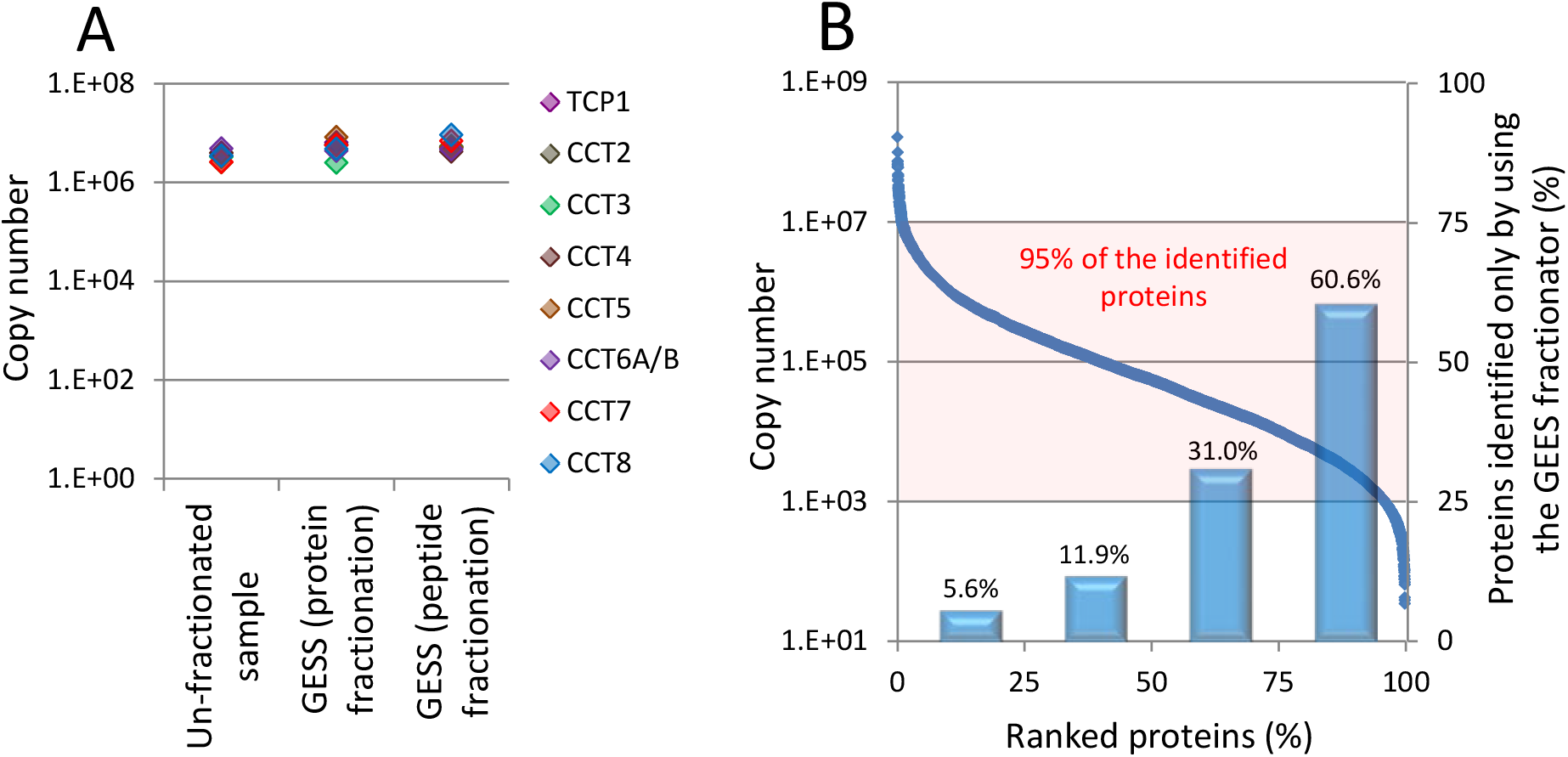
Calculated copy number of the CaSki cell proteins using the “proteomic ruler” approach [52]. (A): copy number of the TRiC chaperone complex (T-complex protein 1) members. TCP1, CCT2, CCT3, CCT4, CCT5, CCT6A/B, CCT7 and CCT8: T-complex protein 1 subunits alpha, beta, gamma, delta, epsilon, zeta, eta and theta respectively. (B): copy number of the identified proteins with at least two unique assigned peptides. Proteins were ranked according to their abundances from highest to lowest copy number in 25% ranged groups. Bars indicated the percent of the proteins identified only when the GESS fractionator was included in the sample processing. Shadowed area indicates the portion of the identified proteome (95%) with copy number ranging from 10^3^-10^7^.

The copy number of the identified proteins ranged 7 order of magnitude (i. e. 10^1^-10^8^ copy number), although the 95% of them are within 4 orders of magnitude (i. e. 10^3^-10^7^ copy number) (Figure 6B). The 94.4% of the most abundant proteins (25% identified proteins with the highest estimated copy number) were identified either in the un-fractionated or in the GEES-fractionated samples. However, the 60% of the identified proteins with the lowest estimated copy number (lower 25%) were identified only when the GEES fractionator was used as the pre-fractionation method (Figure 6B).

## Concluding remarks

In spite of the impressive current progress in mass spectrometry, decreasing sample complexity, by means of protein fractionation, substantially increases the possibilities of detecting very low abundant proteins and sub-stoichiometric post-translational modifications of proteins. The SDS-PAGE technique has an unparalleled resolution for protein separation, covering virtually the entire wide range of protein hydrophobicity, charge and molecular mass. At the same time, the performance of SDS-free PAGE is comparable to other RP-orthogonal electrophoretic or chromatographic techniques for peptide fractionation. Therefore, the development of new equipments that would overcome the well-known drawbacks of the PAGE techniques (i.e. low protein and peptide recovery) could be extremely useful according to the state-of-the-art of proteomics. The GEES fractionator was designed to solve the current limitations of the electrophoretic slab-gel systems.

The reproducibility, protein recovery and loading capacity of the prototype presented here is suitable for proteomic applications. Combined with FASP methods, the use of the GEES fractionator increased the number of identified proteins in Bottom-up proteomics by means of both, protein and peptide fractionation. The in-depth description of the CaSki cells proteome validates the potential of the GEES-FASP approach.

According to the high-quality separation of the protein fractions recovered in-solution, it is conceivable that the GEES fractionator can be used in Top-Down proteomics. Additionally, the GEES fractionator is not limited to PAGE techniques and proteomic studies, as other electrophoretic techniques based on different polymers, could also be potentially adapted. Agarose gel-based electrophoresis for DNA or RNA separation, native electrophoresis for the fractionation of protein complexes, as well as the electrophoretic separation of charged lipids, glycosides or metabolites, among others, might be evaluated in the future.

## Supporting information

Supplementary Information III

## Acknowledgements

The authors thank to Katharina Zettl for technical support as well as to Nayade Beltran and Walter Ramirez for the supplemental video recording and editing.

## Supplementary Information

Supplementary Information I: Gel Electrophoresis/Electroelution Sorting Fractionator functioning (video).

Supplementary Information II: Protein recovery of the SDS-PAGE based GEES fractionation.

Supplementary Information III: Proteins identified per fraction in the CaSki cell extract by combining the SDS-PAGE based GEES protein fractionation and the MED-FASP method (sheets 1-12: Fractions 1-12 respectively).

Supplementary Information IV: Proteins identified in the un-fractionated CaSki cell extract by using the MED-FASP method.

Supplementary Information V: Proteins identified per fraction in the CaSki cell extract by combining the FASP method and the SDS-free PAGE based GEES peptide fractionation (sheets 1-12: Fractions 1-12 respectively).

