## Supplementary Information II for "Gel Electrophoresis/Electroelution Sorting Fractionator combined with Filter Aided Sample preparation (FASP) for deep proteomic analysis"

**Supplementary information II**: Protein recovery of the SDS-PAGE based GEES fractionation.

*Sample preparation*: Bio-Rad Molecular Weight Marker solution as well as CaSki protein extraction were incubated 5 min at 95˚C in electrophoresis sample buffer (SDS 1%, β-mercaptoetanol 2.5%, glycerol 12.5%, Tris/HCl 62.5 mM, pH 6.8). CaSki cells extraction was sonicated. For cysteine modification, acrylamide was added to 6.25% (w/v) and the samples were incubated 1h at 25˚C. Insoluble material was removed by centrifugation at 16,000 x g, 20˚C for 20 min.

For the Bio-Rad Low Molecular Weight Marker, protein recovery was estimated by the Abs_280_ ratio of all the summed fractions to the un-fractionated sample. After the SDS-PAGE based GEES protein fractionation, the recovered fractions were combined and the Abs (280 nm) was measured and compared with the Abs (280 nm) of the corresponding un-fractionated sample. The experiment was carried out by triplicate at different days (figure 1).

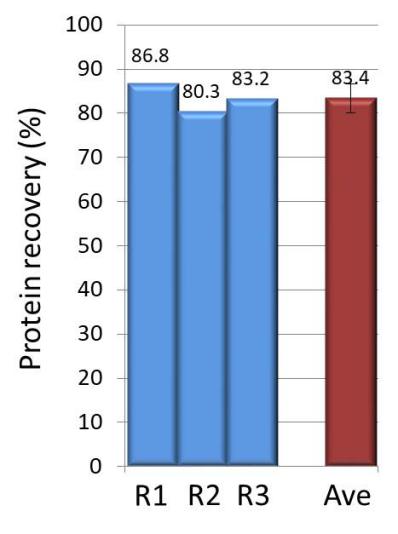

Figure 1: Protein recovery of the SDS-PAGE based GEES fractionation using the Bio-Rad Low Molecular Weight Marker. The experiment was carried out by triplicate. R1-3: Replicate 1-3. Ave: Protein recovery average.

For the CaSki cell extract, protein of the un-fractionated sample (150 µg) as well as the GEES fractions were determined by a 96-Well-Plate-Based Tryptophan Fluorescent Assay as described in [^[1]^]. Protein recovery was calculated as the total protein recovered in all fractions respect to 150 µg (figure 2).

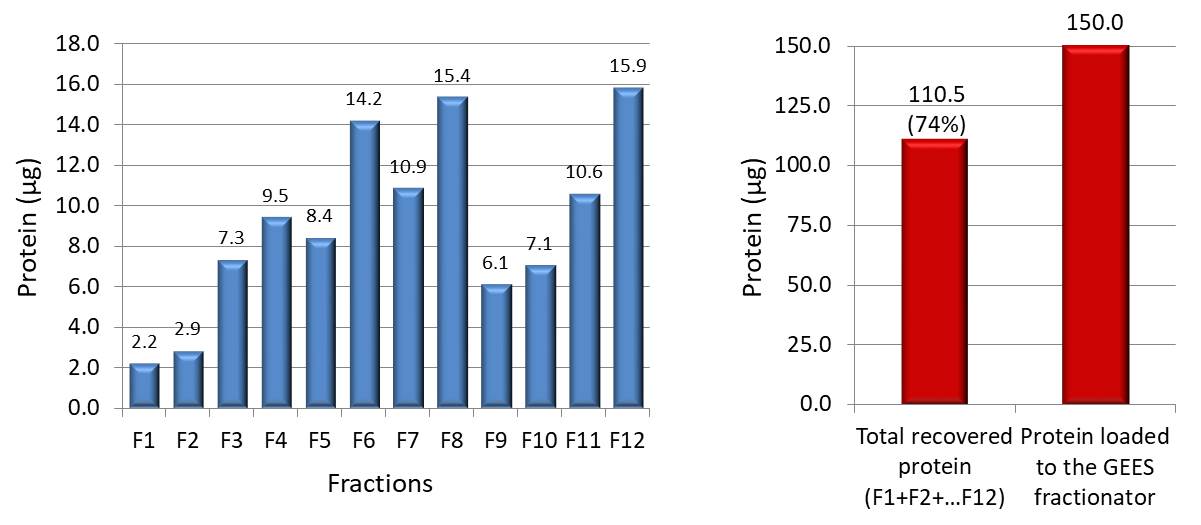

Figure 2: Protein recovery of the SDS-PAGE based GEES fractionation using the CaSki protein extract. Protein recovery was calculated as the total recovered protein of all fractions respect the amount of protein loaded to the GEES fractionator.

1. Jacek R. Wiśniewski, and Fabienne Z. Gaugaz. Fast and Sensitive Total Protein and Peptide Assays for Proteomic Analysis. Anal. Chem. 2015, 87, 4110−4116.
